# Mutations in yeast are deleterious on average regardless of the degree of adaptation to the testing environment

**DOI:** 10.1101/2024.01.09.574908

**Authors:** Kevin Bao, Brant R. Strayer, Neil P. Braker, Alexandra A. Chan, Nathaniel P. Sharp

## Abstract

The role of spontaneous mutations in evolution depends on the distribution of their effects on fitness. Despite a general consensus that new mutations are deleterious on average, a handful of mutation accumulation experiments in diverse organisms instead suggest that of beneficial and deleterious mutations can have comparable fitness impacts, i.e., the product of their respective rates and effects can be roughly equal. We currently lack a general framework for predicting when such a pattern will occur. One idea is that beneficial mutations will be more evident in genotypes that are not well adapted to the testing environment. We tested this prediction experimentally in the laboratory yeast *Saccharomyces cerevisiae* by allowing nine replicate populations to adapt to novel environments with complex sets of stressors. After >1000 asexual generations interspersed with 41 rounds of sexual reproduction, we assessed the mean effect of induced mutations on yeast growth in both the environment to which they had been adapting and the alternative novel environment. The mutations were deleterious on average, with the severity depending on the testing environment. However, we find no evidence that the adaptive match between genotype and environment is predictive of mutational fitness effects.

## Introduction

Many mutations are deleterious or neutral, but some serve as the basis for novelty and adaptation. Predicting the distribution of fitness effects of new mutations (DFE) is an important challenge in evolutionary genetics (1,2). Methods are being explored to experimentally characterize the fitness effects of many individual alleles (3–5), and to predict mutational fitness effects based on molecular data (6–8). While these are exciting prospects, it is not clear that we can currently predict basic elements of the DFE, even in model organisms. A key method for studying mutational effects is mutation accumulation (MA), where selection is rendered largely ineffective in replicate lineages by repeated bottlenecking to minimize the effective population size (9–11). Most such studies show a decline in mean fitness under MA, indicating that beneficial mutations must have a much smaller net impact than deleterious mutations, i.e., they occur less frequently, have weaker effects, or both. However, fitness decline is not always observed in MA experiments, even when other evidence confirms that un-selected mutations are accumulating (reviewed in (12)). The simplest interpretation of this pattern is that beneficial mutations were relatively common, with a net impact on fitness similar to that of deleterious mutations (5,13,14). We should ideally be able to predict whether beneficial mutations will be common or scarce, in order to understand the role of mutation in evolution.

An attractive explanation for the presence of inconsistent consequences of MA has to do with the extent of prior adaptation. Intuitively, a genotype that is well adapted to a given environment would have little beneficial mutational variation available for further adaptation, relative to a poorly-adapted genotype in the same environment––minimal adaptation to the testing environment could perhaps account for the lack of mean fitness decline observed in some MA experiments. Such an outcome might be more likely for organisms that are preserved in freezers or seed banks prior to experimentation (e.g., yeast, plant seeds), as opposed to those maintained continually in lab environments (e.g. *Drosophila*). Using a “fitness landscape” analogy, a population on a fitness peak would have nowhere to go but down, but one further from the peak would have opportunities for fitness improvement. Such models have been explored extensively, evolving from ideas by Fisher as a part of the eponymous Geometric Model (15) to more detailed mathematical treatments (16–20). Extensions of Fisher’s model assuming a Gaussian fitness landscape predict that the current “adaptedness” of a genotype influences the variance of mutational fitness effects but not the average effect; this is because while relatively more beneficial mutations are available to a maladapted genotype, this is offset by an increase in the severity of deleterious mutations (17). Experimental tests of these predictions have yielded somewhat mixed results (21,22), which we review briefly below (see also (12,23)).

To study the role of adaptation in shaping the DFE, Wang et al. (21) studied the fitness effects of 36 gene disruption alleles back-crossed into *Drosophila melanogaster* genotypes that had previously adapted to one of two alternative environments (cadmium- and salt-enriched larval media diets), when measured in each environment. They found that these alleles were generally deleterious regardless of whether the genetic background was well-adapted to the assay environment or not, and that the environment per se had the largest effect on the mean selection coefficient, rather than evolutionary history: selection against gene disruptions was stronger on average when assessed in the salt environment regardless of genetic background. Genetic background and adaptedness were also found to have no strong effect on the variance in selection coefficients, whereas environment did have an effect on the variance. In another study (22), *Arabidopsis thaliana* lines from the extremes of its natural range (southern France and central Sweden) were used to test the predictions of fitness landscape theory. Founder plants collected from these two range extremes were cultivated for 7-10 generations under MA conditions. They then assessed the change in fitness in these mutant lines in both the founder’s original environment (“home”) and the other environment (“away”). MA lines with a fitness difference from the ancestor showed a nearly universal trend of fitness decline. MA lines also performed worse in the novel sites than in those sites where their founding genotype originated, contradicting the prediction that there would be no difference in the mean fitness effects of new mutations. The authors concluded that the mean fitness effect of mutations depends more on the environment than on adaptedness, similar to the Wang et al. (21) study. While variance among lines was positively correlated with how stressful the environment was (and away sites were universally more stressful than home sites for the mutants), among-line variance accounted for very little of the overall variation (22).

While compelling, these studies naturally have limitations. Wang et al. (21) studied specific gene disruption mutations on a single chromosome. While this allowed for the effects of specific, individual mutations to be assessed, spontaneous mutations have a more diverse molecular spectrum, affecting both genic and non-coding regions, which can both have effects on fitness (1,24). In another study, Weng et al. (22) relied on natural populations of a species with well-documented patterns of local adaptation. The use of natural variation brings with it all the challenges presented by any wild population like standing genetic variation, demographic structure, and an uncertain evolutionary history. Further complicating the picture, several different fitness metrics were used, which produced somewhat inconsistent results. Finally, although they did find increased variance in the more stressful environment as predicted by the Martin and Lenormand model (17), this could have been due to standing genetic variation that only influenced fitness in more stressful environments (25). These studies in very different organisms reflect the challenges of testing the predictions of the fitness landscape model, perhaps explaining why experiments of this kind remain scarce.

Our goal was to investigate how new mutations affect fitness given alternative histories of adaptation in the budding yeast *Saccharomyces cerevisiae*. In this species there are examples of MA apparently resulting in little mean fitness decline (14,26), suggesting that the average effect of mutations may be influenced by the environment or genetic background. We followed previous studies by using a reciprocal transplant-style design, which allows effects of adaptation to be distinguished from effects of the environment. We allowed diploid *S. cerevisiae* populations to adapt to novel environments that each contained multiple stressors, with regular opportunities for sexual reproduction within populations to facilitate adaptation (27). We chose to create relatively complex environments to minimize the possibility that adaptation would involve only a single genetic pathway (28,29), which might have idiosyncratic interactions with new mutations. After generating these “locally adapted” populations, with highest fitness in their “home” environments, we measured the fitness consequences of induced mutations in both “home” and “away” environments, for each population. We found that mutations had deleterious effects on average, regardless of whether the genetic background was well- or poorly-adapted to the testing environment.

## Methods

### Strains, growth measures and media

We obtained the haploid strain FM1282 from C. Hittinger, which was derived from BY4724 and modified to express green fluorescent protein (30). By inducing mating-type switching and mating we generated a homozygous diploid version of this strain with genotype *ho ura3*Δ *lys2*Δ PTDH3-yEGFP-TCYC1 to use in our experiments; we preserved this strain at –80 C, and hereafter we refer to it as the “ancestral” strain. We cultured the yeast at 30 C with continuous shaking applied to the liquid cultures. We used growth rate assays in several stages of our study, which we conducted in a BioTek Epoch II plate reader, taking regular measurements of optical density (OD) at 600 nm with incubation and shaking. We calculated maximum growth rate in each assay culture by fitting a spline to the log of OD over time using the *loess* function in *R* with degree = 1 and span = 0.2 and finding the maximum slope of this spline across 99 equally-spaced time points.

Our first goal was to generate three liquid-media environments that resulted in moderately reduced yeast growth rates. Based on literature searches we identified several substances known to affect growth rate and tested their effects when added individually to liquid yeast-peptone-dextrose (YPD) media to find appropriate concentration ranges. In order to generate “complex” environments we combined groups of four substances and performed additional growth rate tests. We chose combinations of substances haphazardly, but we sought to make each environment distinct, and excluded any combinations found to precipitate out of solution. The final media environments used in our experiments contained the following substances dissolved in YPD. We called these environments EnvA, EnvB and EnvC; *EnvA*: boric acid, caffeine, nickel chloride and sodium fluoride; *EnvB*: acetic acid, acetaminophen, lithium chloride and sodium chloride; *EnvC*: chromium potassium sulfate, cupric sulfate, sodium benzoate and zinc chloride. Recipes and concentrations for each environment are given in Table S1. These three environments all produced similar reductions in growth rate of the ancestral strain, relative to its growth rate in YPD (Fig. 1).

**Figure 1.**
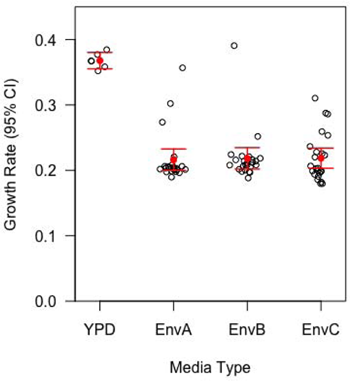
Initial media environments reduced the growth rate of laboratory yeast. All three of the newly formulated environments had a significant impact on growth rate when compared to standard laboratory YPD (t-tests; EnvA: *P* = 2.566 × 10^-15^; EnvB: *P* = 3.387 × 10^-15^; EnvC: *P* = 2.473 × 10^-15^). The three novel environments all reduced growth to a similar extent. Growth rate refers to the rate of change of OD per hour.

### Adaptation

For each of the three environments we initiated three replicate yeast populations, each growing in 3 mL of liquid media. We passaged each population three times a week (every 2-3 days) by inoculating 3 mL of new media with 30 μL of the previous culture, i.e., a 1:100 dilution. We refer to populations that were serially passaged through these environments as A-evolved, B-evolved, and C-evolved, respectively. At the end of each week, we sampled yeast from each population and assessed their growth rate in their assigned media, alongside the ancestral strain (see below). Relative fitness is calculated as the difference in growth rate between the ancestor and the evolving populations. This allowed us to monitor the relative fitness of the adapting populations on a regular basis. We froze samples from each population in 15% glycerol at regular intervals throughout this phase as well as at the end of the adaptation. These procedures are depicted in Fig. 2A.

**Figure 2:**
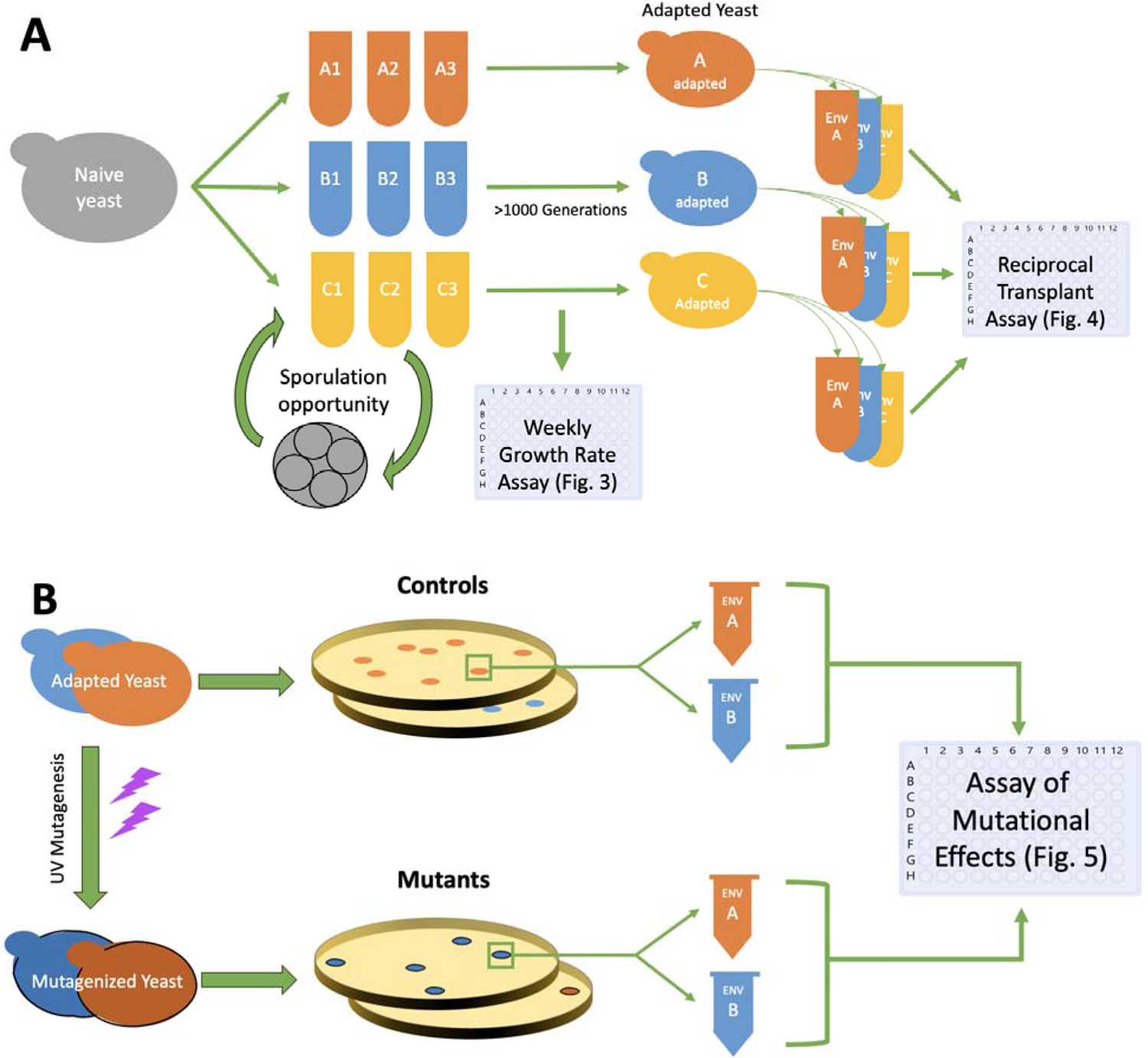
Diagrammatic representation of the experimental protocol. Panel A depicts the experimental steps that were undertaken during the “adaptation” phase of the experiment where we took populations of yeast and subjected them to challenging environments for many generations, with periodic opportunities for sexual reproduction (sporulation). The results of this part of the experiment are shown in Figures 2 and 3. Panel B depicts the steps taken to mutagenize yeast and then measure their fitness in both familiar and novel environments. The results of these experiments are shown in Figure 5.

Every third week we allowed the adapting populations to undergo sexual reproduction within their own population (independent replicate populations were not mixed). We inoculated 3 mL of pre-sporulation medium (1% potassium acetate, 1% yeast extract, 2% peptone), with 30 μL of saturated culture for each replicate population and allowed the cultures to grow for 18-24 h with shaking at 30 C. We then centrifuged these cultures, resuspended the cells in 3 mL sporulation medium (1% potassium acetate) supplemented with amino acids (0.01% lysine and uracil), and incubated them at room temperature on a rotor for 72-120 h. These conditions encourage sporulation but limit the opportunity for asexual growth. We then examined these cultures for the presence of tetrads, but used all cells to inoculate 3 mL of new media, allowing asexual growth to resume in the appropriate media environment for each replicate population. Note that this procedure allowed for sexual reproduction to take place but did not enforce it, as non-sporulated cells were not eliminated. We assume that haploid spores germinated upon returning to nutritive media, and mated to re-form diploid cells, but we also confirmed ploidy at the end of the adaptation phase (see below). During the weeks when sporulation was permitted, we did not freeze any of the adapting populations or conduct fitness assays. The sporulation media did not include our stressful additives, and so we do not include the sporulation periods when calculating the total time spent adapting to the novel media environments. In total, the experimental populations spent 357 days growing asexually in their respective media. Given the number of 100-fold dilutions, 160, there must have been at least 160*log*_2_(100) = 1063 rounds of asexual cell division over the course of the experiment. At ten timepoints throughout the adaptation phase we performed cell counts to estimate the size of the experimental populations just prior to passaging. For the cultures we ultimately used in tests of mutation effects, the average population size at passaging was 7.7 × 10^8^ cells per mL (SE 4.2 × 10^7^, *n* = 106), indicating effective population sizes of at least 7.7 × 10^6^, i.e., the estimated population size immediately following each dilution.

### Growth rate assays during adaptation

Throughout the adaptation phase of the experiment, we conducted regular assays to track the growth rate of the evolving populations in their respective media environments relative to the ancestor. Prior to each assay, we revived the ancestor strain, which had not undergone any adaptation, from frozen stock and grew it in YPD for two days. We set up a 96 well plate with a stratified treatment arrangement; each plate held 21 wells each of A-evolved, B-evolved, and C-evolved in their respective media types, 9 wells each of the ancestor growing in EnvA, EnvB, and EnvC for comparison, and two “blank” wells per media type, not inoculated with yeast, to detect contamination. We diluted each yeast culture 500-fold to initiate the growth assay, and obtained OD readings every 15 min for 30 h.

### Growth rate assays following adaptation

Following the adaptation phase of the experiment we assessed whether each population grew more rapidly in its “home” environment than in the alternative, “away” environments. We also included the ancestral strain in these assays for comparison (Fig. 2A). We designed these assays to be similar to the transplantation assays used in other studies to verify that local adaptation had occurred (22), i.e., that the best performing populations in a given environment were those with a history of adaptation to that environment. We conducted these assays using the same methodology and setup as the weekly fitness assays conducted during the adaptation phase of the experiment except that we collected data for 48 h rather than 30 h to account for the possibly slowed growth rates in novel environments. We tested each adapting population in each of the three environments, effectively “transplanting” lines from familiar “home” environments they had adapted to (e.g., B-adapted populations in Env B) to novel “away” environments(e.g., B-adapted populations in EnvA). We additionally tested the ancestral strain in each environment. As described below (see Results), we found that the expected signature of local adaptation was absent for the C-evolved populations, and so we excluded them from the remainder of the experiment. We repeated this assay using only A-evolved and B-evolved populations in EnvA and EnvB, testing six replicates of each evolved population in each environment, and four replicates per environment for the ancestral strain. Relative growth rate was calculated as the difference in growth rate between an evolved population in a given environment and that of the ancestor in the same environment.

### Mutagenesis and mutant fitness assay

We counted the number of cells in saturated samples from each population grown in their home environments, and used this number to plate 100–200 single cells on each of three YPD agar plates. We then placed two plates in a biosafety cabinet, approximately 84 cm from a germicidal UVC bulb, and irradiated them for 10 seconds with their lids removed, with the remaining plate serving as a control. Following 2 d of incubation we counted colonies on irradiated and non-irradiated plates, and observed an expected 40-70% reduction in colony counts for all irradiated plates (31), representing a significant effect of UV on cell survival. This is taken to indicate that mutagenesis was effective in generating non-lethal mutations in the surviving cells. For each strain we chose one irradiated plate at random and picked several colonies from this plate and from the the non-irradiated plate using sterile toothpicks. To prevent bias, we picked colonies from the center of the plate moving outwards. We suspended each colony in sorbitol before dividing the volume into two 1.5 mL centrifuge tubes, followed by a 10-fold dilution in sorbitol.

These colonies each represent a mutant line and will be referred to as “mutants” and mutant or mutagenized “genotypes.” We centrifuged these tubes, removed the sorbitol, and resuspended the cells from one tube in EnvA and the other in EnvB. In this way we were able to measure the growth rate of each mutant genotype, as well as the non-mutagenized control, in both EnvA and EnvB. We measured growth rates using the same procedures as the fitness and transplantation assays described above, and collected data for 48 h. In total, we measured the growth rate of 162 mutagenized genotypes and 114 non-mutagenized genotypes in both EnvA and EnvB, for a total of 552 measurements. Here, relative growth rate is the difference in growth rate between a mutant in a given environment and a non-mutagenized control with the same evolutionary history in the same environment.

A class of mutation that may occur in irradiated yeast is respiratory deficiency, resulting in the “petite” phenotype (32,33). This could be problematic for our fitness assays because if this type of mutation is common it could have an outsized impact on fitness relative to other mutations. To determine how frequently our mutagenesis protocol produced mutants with the petite phenotype we repeated the UV mutagenesis procedure and selected mutant colonies following the same procedure used for the mutant fitness assays. We then patched these mutants onto YPD agar plates alongside non-mutagenized controls. After 3 d of growth at 30 C, we replica plated onto YPG (yeast peptone glycerol) agar plates, a non-fermentable medium on which petite strains cannot grow, allowing us to determine the frequency of petites.

### Flow cytometry

While we allowed for sexual reproduction (sporulation) periodically throughout the adaptation phase of our experiment, we expected the resulting haploid spores to rapidly germinate and mate with one another upon encountering nutritive media, thereby restoring the diploid state. However, unicellular fungi have been known to change ploidy during adaptation (34,35), and so we confirmed ploidy using flow cytometry for each experimental population following adaptation, using known haploid and known diploid versions of the ancestral strain for comparison.

Following overnight growth, we combined 200 µl of each culture with 800 µl pure water, pelleted cells, and gently resuspended in 1 mL cold 70% ethanol. After 1 h incubation at room temperature we pelleted and washed cells twice with 1 mL sodium citrate (50 mM, pH 7), then added 25 µL of RNAse A (10 mg/mL) and incubated overnight at 37 C. We then pelleted and resuspended cells in sodium citrate and added 30 µL of 50 µM SYTOX green nucleic acid stain (Invitrogen S7020). Finally, we incubated samples in the dark overnight and performed analysis of particle fluorescence and size on an Attune NxT V6 flow cytometer (ThermoFisher). We removed particles with extreme size or shape from the dataset, and visually compared the fluorescence profiles of known haploids and diploids with those of our experimental populations. These analyses confirmed that all nine of our experimental populations consisted of cells in the diploid state by the end of the adaptation phase (Fig. S1).

## Results

### Adaptation

Throughout the adaptation phase of the experiment, we visually confirmed the presence of tetrads in all adapting populations during each week of sporulation, indicating that some propensity to undergo sexual reproduction was maintained. After 372 days of asexual growth (>1000 generations) in the novel media environments (not including time spent in sporulation media), all populations of A-evolved, B-evolved, and C-evolved showed a significant improvement in fitness when compared to the ancestral strain grown in the same environment (Fig. 3): in a mixed-effect model of growth rate accounting for random effects of population, environment, and assay block, we find a highly significant interaction between evolution status (adapting or ancestral), and time (days of evolution) (likelihood ratio test (LRT) χ^2^ = 125.13, *P* < 10^-15^). A random effect of assay plate location did not have a significant impact on our results when factored into the linear models (LRT: χ^2^ = 5.20, *P* = 0.074), and was omitted. To summarize, the experimental populations all performed better than the unadapted ancestors in their respective media types.

**Figure 3.**
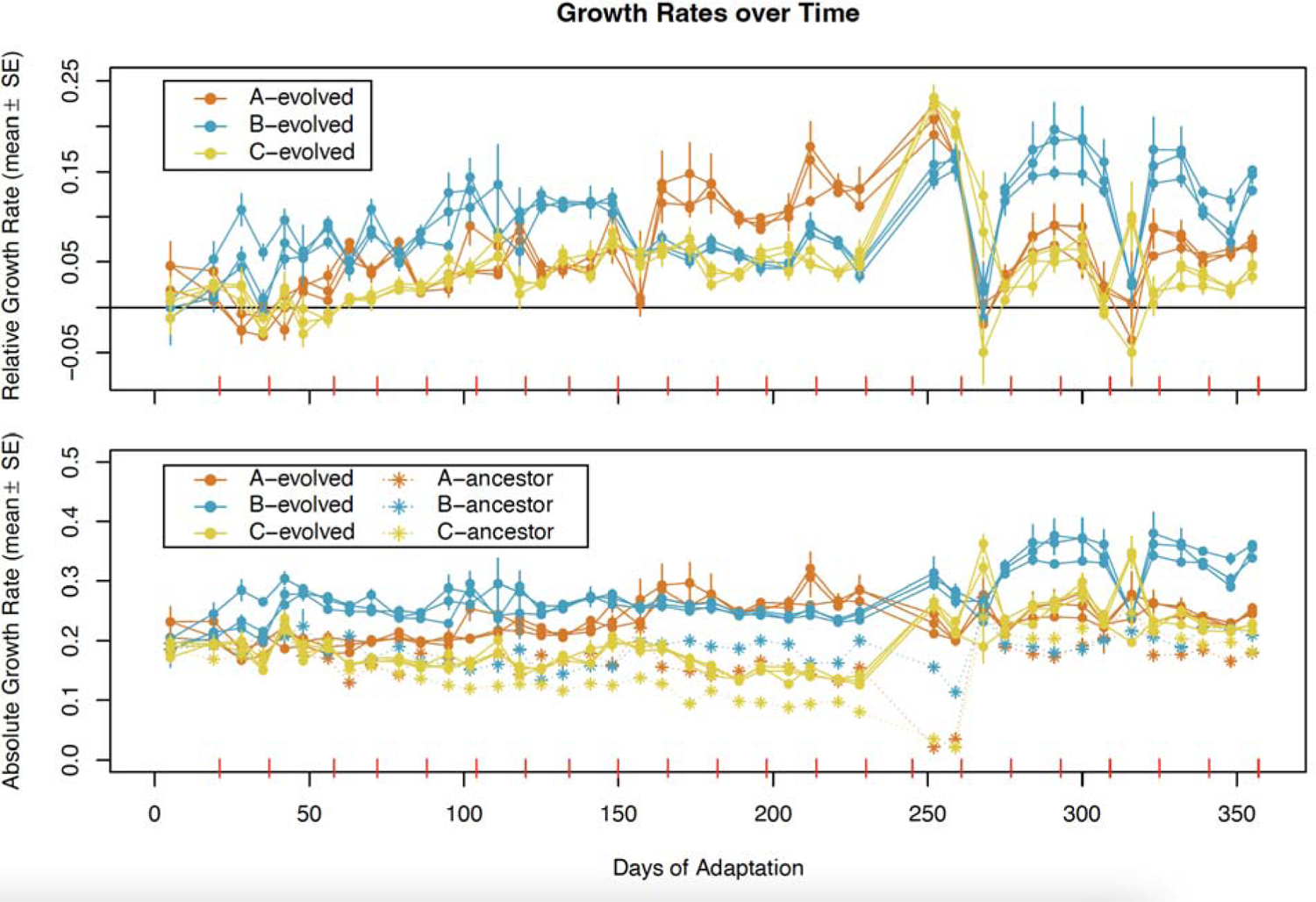
Growth in novel environments improved over time. The top panel shows growth rate of the evolving populations relative to ancestors while the bottom shows the absolute growth rate of ancestors and evolving lines. Red tick marks on the x-axes indicate sporulation opportunities. These were not counted toward total days of adaptation because sporulation medium was permissive. By the end of the experiment, all evolving populations showed substantially improved growth when compared to the ancestor.

### Transplantation assay

We performed a transplantation assay to identify signatures of “local adaptation”. Our initial assay tested each permutation of evolved populations and ancestral, unadapted yeast with each of the environments EnvA, EnvB, and EnvC (Fig. 4A). Using a mixed-effect model of growth rate with random effects of population and assay block we found a highly significant interaction effect between evolutionary history (i.e., the environment in which a population evolved) and the test environment (excluding the ancestor; LRT, χ^2^ = 67.892, *P* < 1 × 10^-15^). We then analyzed the effect of evolutionary history in each test environment separately, and detected significant effects of evolutionary history in EnvA (LRT, χ^2^ = 3.8743, *P* = 0.049) and EnvB (LRT, χ^2^ = 6.97, *P* = 0.0083). In EnvC we did not detect such an effect (LRT, χ^2^= 0.7511, *P* = 0.39), meaning that populations that had evolved in EnvC did not outperform other evolving populations in that environment (Fig. 4A). Notably, populations evolving in EnvC also showed the weakest improvement in relative growth rate in our weekly fitness assays (Fig. 3).

**Figure 4.**
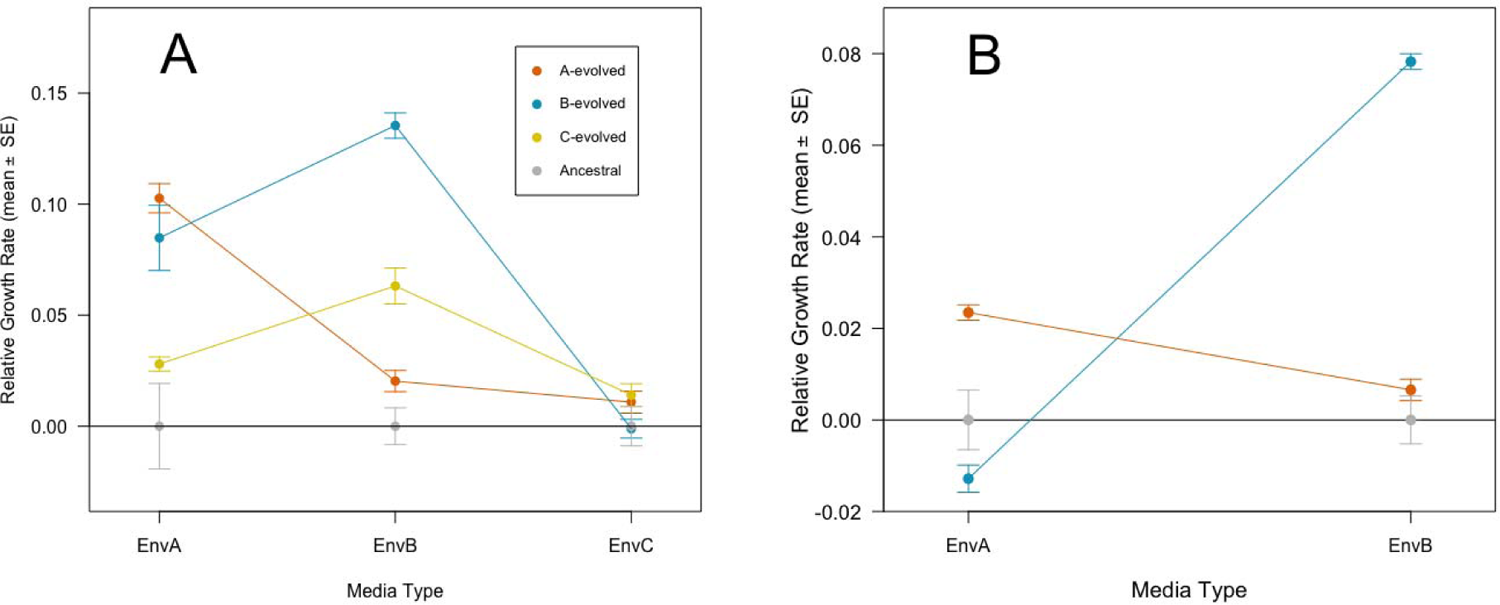
Transplantation assays reveal signatures of local adaptation. Shown here are the relative growth rates compared to ancestors in our first (A) and second (B) transplantation assays. The initial assay (A) showed the expected signature of adaptation was absent for C-evolved lines, so we repeated the assay (B) with only A-evolved and B-evolved lines and their respective environments. In panel (B) we can see that yeast perform best in the environments to which they had been adapting. Ancestors are included to show the standard error for unadapted lines. The data plotted here are also available in Table S2.

Because the C-evolved populations failed to show a signature of local adaptation, we repeated the transplantation assay with only A-evolved and B-evolved populations and EnvA and EnvB (Fig. 4B). Repeating our analyses with mixed effect linear models with replicate population and plate as random effects, we again found a highly significant genotype by environment interaction effect on growth rate (excluding the ancestor; LRT, χ^2^= 298.21, *P* < 10^-15^). The testing environment had a significant effect on growth rate for both A-evolved (LRT, χ^2^ = 34.893, *P* < 10^-8^) and B-evolved (LRT, χ^2^ = 225.02, *P* < 10^-15^) populations, where growth was highest in the environment to which each population had adapted (Fig. 4B). We reanalyzed the data using the *emmeans* package in *R* to obtain post-hoc contrasts accounting for multiple testing, pooling the replicates of the adapting populations within treatment groups after determining the replicate populations did not differ statistically (all χ^2^ < 3.12, all *P* > 0.077). Contrasts showed that each adapted population performed better than unadapted ancestors in EnvA (*t* = 5.170, *P* < 0.0001) and EnvB (*t* = 10.438, *P* = 0.0067). We then asked if the evolved populations performed differently from the ancestor in non-familiar environments (e.g. A-evolved grown in EnvB). The adapted populations did not perform significantly differently than the ancestors in these comparisons (A-evolved vs. ancestor in EnvB: *t* = 1.445, *P* = 0.47; B-evolved vs. ancestor in EnvA: *t* = –1.713 *P* = 0.45). This shows that adaptation to a given environment did not result in significantly improved fitness in another environment with a different set of stressors. This can also be taken as an indication that general lab adaptation played little to no role in the fitness of the adapted populations.

### Mutant fitness

Our assessment of respiratory deficiency in mutagenized cells indicated that this type of mutation was not widespread: about 2% of mutagenized cells formed petite colonies (8/407 colonies tested), and this was not significantly higher than the petite frequency in the non-mutagenized treatment (0/141 colonies tested; Fisher’s exact test, *P* = 0.212). Among the mutagenized colonies we used for our fitness assay, we would therefore expect only two or three petites, and so this type of mutation should not have undue influence on our results.

We measured the fitness of mutagenized genotypes and their non-mutagenized counterparts. Each genotype was measured twice: once in EnvA and once in EnvB. To compare groups we used bootstrapping with 10 000 replicates, retaining the paired nature of the data, i.e., the fact that each genotype was measured both EnvA and EnvB. First, we observed that the mutagenized genotypes had significantly lower growth rates than non-mutagenized genotypes, averaging over adaptive history and testing environments (bootstrap *P* < 2 × 10^-4^), demonstrating that mutagenesis was effective in generating (predominantly deleterious) mutations. The average magnitude of fitness reduction caused by mutagenesis was equivalent to approximately 6458 generations of spontaneous mutation accumulation, using data from the wild-type diploid MA lines of Sharp et al. (26) as a standard. For the A-evolved populations, mutagenesis did not significantly reduce average growth rate in EnvA (bootstrap *P* = 0.63) but did so in EnvB (bootstrap *P* = 0.0124). For the B-evolved population, mutagenesis resulted in reduced average growth rates in both EnvA (bootstrap *P* < 2 × 10^-4^) and EnvB (bootstrap *P* < 2 × 10^-4^) (Fig. 5).

**Figure 5.**
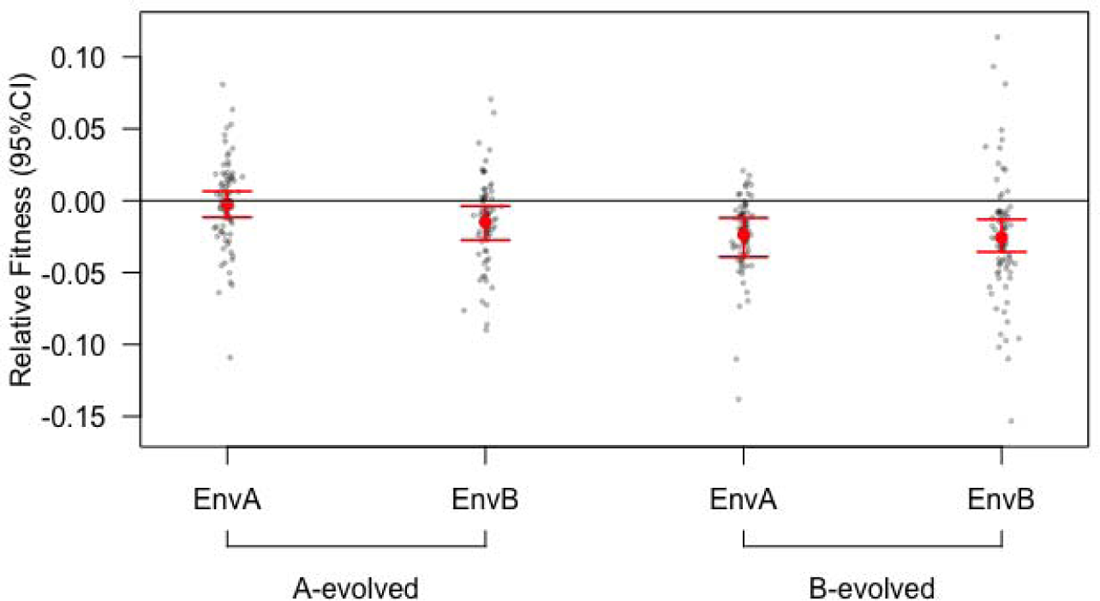
The effect of new mutations given alternative histories of adaptation. Red dots indicate the mean relative fitness of the group of mutants in a given environment while the smaller black dots indicate the individual mutants. Relative fitness here represents the difference in growth rates between a given mutant in a given environment and a non-mutagneized control with the same adaptive history (originally from the same evolving population) in the same environment. 95% confidence intervals were obtained from 10 000 bootstrap replicates. Mutations were consistently deleterious on average regardless of adaptive history. Exact values can be found in Table S3.

Our primary interest was in whether the average effect of mutations depended on the degree of adaptation to the testing environment. If the mean fitness effect of mutations were to become more beneficial in maladapted genotypes, we would expect to find a higher mean relative fitness in mutants assessed in “away” environments than in “home” environments. Our results do not show this pattern. Indeed, mutations were more deleterious on average in the away environment than in the home environment for A-evolved population mutants (bootstrap *P* = 0.0466). For the B-evolved case, mutational effects did not differ between environments (bootstrap *P* = 0.86). Averaging across evolutionary environments, there was no evidence that adaptedness influenced the average effect of mutations (bootstrap *P* = 0.30). Given this lack of a genotype-by-environment interaction effect, we can test for main effects of each factor. On average, the fitness effects of mutations were more deleterious in the B-evolved genetic background than the A-evolved genetic background (bootstrap *P* = 0.0132), but we do not find evidence for a difference in mutational effects between EnvA and EnvB (*P* = 0.172). In other words, we detect a main effect of genotype but not environment.

Fitness landscape theory (17) predicts that the variance of fitness effects of mutations should be higher for genotypes further from the fitness optimum. In our experiment, each mutant genotype was only measured once in each environment, so we cannot estimate genetic variances. However, if we assume a constant value for error variance in all treatment groups, then the following comparisons would be informative. We detected no difference in the phenotypic variance between the two environments for the mutants derived from A-evolved populations (*P* = 0.81). Mutants derived from B-evolved populations did show a significant difference in fitness variance, but it was higher in the home environment than in the away environment (*P* < 2 × 10^-4^) (See Fig. S2). On its face, this is inconsistent with the prediction that variance should be higher for maladapted genotypes, but we cannot rule out differences in error variance among groups.

## Discussion

We investigated ideas about mutational fitness effects in relation to degree of adaptedness by conducting experimental evolution and mutagenesis. By using a fast-growing model organism, we were able to allow for >1000 generations of adaptation to complex, novel environments, while maintaining a high degree of control over environmental conditions. We also allowed for frequent genetic mixing within populations, which may have facilitated the response to selection. We surmise that adaptation to the novel environments most likely involved multiple genetic loci, given the chemical complexity of the environments and the relatively gradual adaptation we observed (Fig. 3). We think this scenario is a better match to what natural populations might experience than, e.g., the presence of a single stressful drug. We did not attempt to identify adaptive alleles in our experimental populations, which could serve to verify that adaptation was multifaceted. However, the theory we set out to test requires only differences in fitness between environments, for which we found strong evidence (Fig. 4). Similarly, our mutagenesis protocol had clear effects on fitness (Fig. 5), but we did not attempt to identify the molecular nature of induced mutations.

We found that random mutations tended to reduce mean fitness, regardless of whether the genetic background on which they arose was well-adapted to the testing environment or not. This finding is not consistent with the idea that the net effect of mutations might be more beneficial in poorly-adapted genotypes. Formal fitness landscape theory also rejects this notion, predicting instead that the increased availability of beneficial mutations in poorly-adapted genotypes is counteracted by an increase in the severity of deleterious mutations, resulting in no effect of adaptedness on the average effect of new mutations. Our results are not perfectly consistent with this prediction either, as we find that mutations in the A-evolved genetic background were more deleterious on average when tested in EnvB versus EnvA. We also do not see greater variance in mutational effects in novel environments, which is predicted by the theory (Fig. S3), though we can examine only phenotypic and not genetic variances.

Our experiment was partly motivated by the finding that MA studies don’t always show a pattern of fitness decline (13,14). We set out to determine if variation in the adaptedness of the initial genotype could help explain this phenomenon. Theoretical investigations of Gaussian fitness landscapes suggest that such an effect should not be expected (17), and an effect of adaptedness has now been rejected by our study of yeast, along with similar studies of new mutations in *Drosophila* (21) and *Arabidopsis* (22); the fact that emprical studies using diverse model organisms and methods for generating mutations all reject a role of adaptedness suggests that this result may be general. At the same time, these experiments also indicate that a simple application of the fitness landscape model may not be sufficient to predict the fitness effects of new mutations, as there are several cases where mutations appear to be more deleterious in maladapted genotypes.

There are limitations to our experiment that could affect our conclusions. While we confirmed that fitness increased over time in our experimental populations (Fig. 3), resulting in a pattern of “local adaptation” (Fig. 4), fitness might have continued to improve if we had allowed even more time for evolution in the novel environments. If the mean effect of new mutations depends on adaptedness, this would presumably be easier to detect when the difference in fitness between adapted and non-adapted genotypes in a given environment is large. One indication that further adaptation may have been possible in our experiment is that the evolved populations still had lower absolute growth rates than the ancestral genotype growing in YPD (compare Fig. 1 and Fig. 3B). Nevertheless, even if the experimental populations had not yet reached new fitness “peaks”, we would still expect the predictions of the fitness landscape model to bear out, just to a lesser degree (36). Given the substantial difference in fitness between our adapted and non-adapted populations, particularly in EnvB (Fig. 4), we should be able to detect an effect of adaptation on mutational fitness effects, if present.

We expect our mutagenesis procedure to generally result in heterozygous mutations, since mutagenized cells did not undergo sex and there was little time for loss-of-heterozygosity events to occur; while this could make it harder to detect the effects of partially recessive mutations (14), selection on heterozygotes is relevant in many natural populations. Additionally, UV mutagenesis is known to create a molecular spectrum of mutations that is somewhat distinct from the spontaneous spectrum (31). Prior experiments of this kind examined X-linked gene dispruption alleles in flies that were hemizygous in males and heterozygous in females (21), or spontaneous mutations in selfing lineages of *Arabidopsis*, which would be largely but not exclusively homozygous by the time of the fitness assays (22).

As in prior studies, we did not find evidence for a role of adaptedness––the interaction between genotype and environment––in determining the average effect of new mutations. Instead, we found that mutations were more deleterious, on average, in B-adapted yeast, regardless of the testing environment. This is in contrast to the prior studies, which both identify the environment, rather than the genetic background, as a main effect (21, 22). A main effect of genetic background could in principle be due to a reduced susceptibility of the A-evolved populations to the mutagenesis.

In conclusion, there appears to be no theoretical or empirical support for the idea that the degree of adaptedness to a given environment has any predictable impact on the average fitness effect of new mutations. On the other hand, attempts to test this idea have identified effects of environment or genotype *per se*, along with other studies (e.g., 37). We can potentially quantify relative adaptedness (or “stress”) in any system, and so it would be convenient if this unifying metric had predictive value for the DFE. Unfortunately, there is more evidence that environment and genetic background, of which there are innumerable potential states, can independently determine how new mutations affect fitness, and so predicting the DFE, or even its average, remains a significant challenge.

## Acknowledgements

Thanks to C. Hittinger for providing yeast strains and to Denise Smith for assistance in the lab.

## Supplementary Information

**Figure S1.**
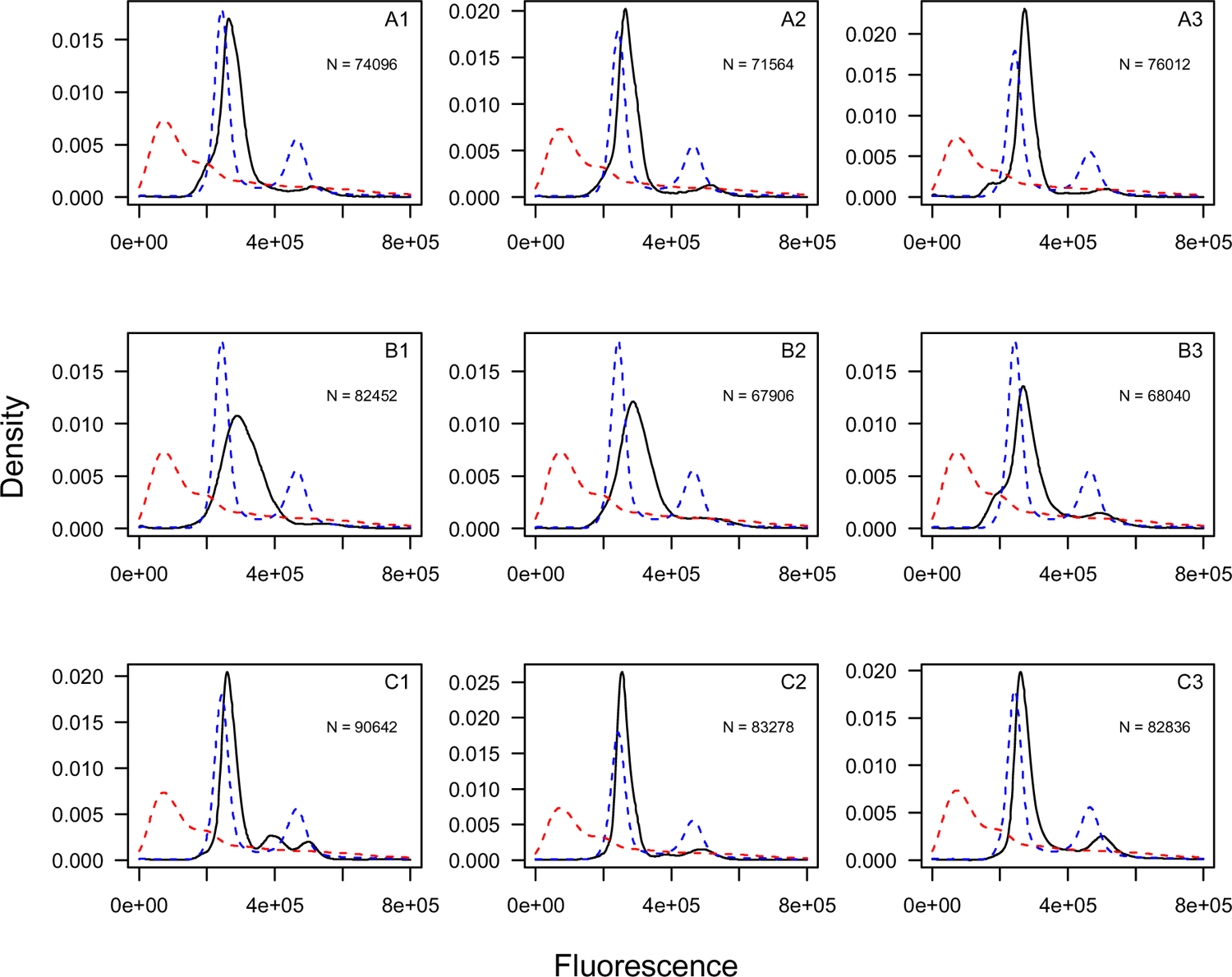
Diploidy was maintained in all experimental populations. Each panel shows the standardized fluorescence density profile for cells from a given experimental population, with the population ID indicated in the top right. Black lines represent the experimental population of interest; dashed lines represent known haploid and diploid strains for comparison (red and blue lines respectively, the same in each panel). Particle numbers are shown in each panel for the experimental populations; known haploids and diploids the plots reflect 44880 and 77041 particles, respectively. In each case, the experimental populations resemble the diploid standard.

**Figure S2.**
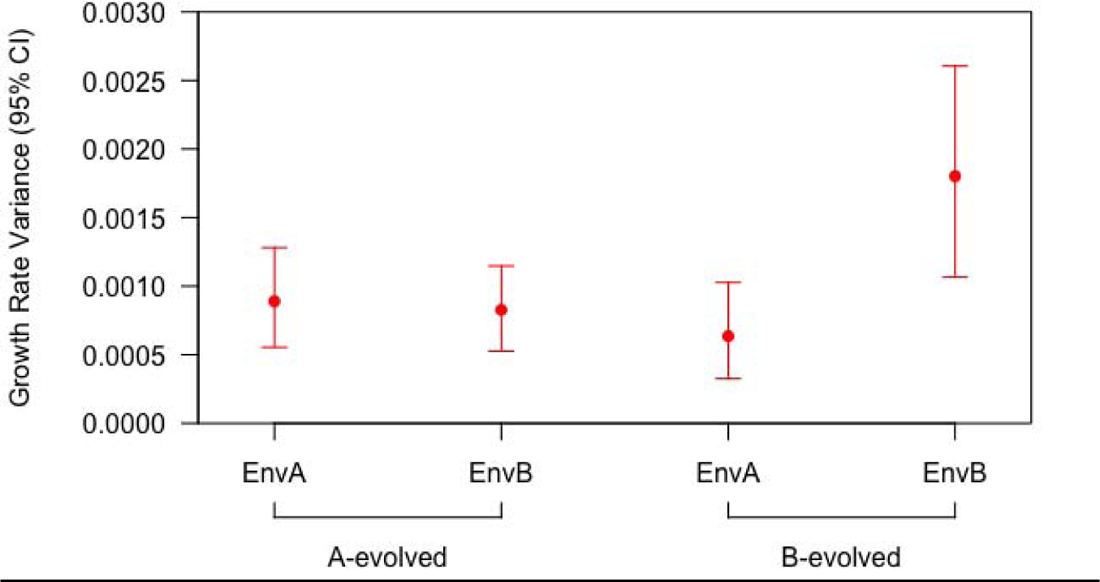
The phenotypic variance of growth rate given alternative histories of adaptation. Dots indicate the variance of each set of mutants in a given environment. The bars indicate the 95% confidence intervals calculated from the same bootstrapped data in Figure 4. It is predicted that mutational variance would be higher when populations were in environments they had not adapted to. Variance did not differ significantly across environments for A-evolved mutants but B-evolved mutants generally had higher variance in EnvB—the opposite of what we would predict. Exact values can be found in Table S2.

**Table S1:** Media recipes. We formulated our environments in a haphazard fashion, aiming to create distinct environments without overlap of stressors between environments or overemphasis of a single stressor type in any given environment. Each environment contains a variety of stressors. We wanted to keep the environments somewhat similar so each one was formulated with 4 unique compounds. Environments were all made by mixing autoclave-sterilized, standard YPD with powdered chemicals and mixed with a magnetic stir bar and gentle heating until fully dissolved. The finalized media with all compounds added was filter sterilized through a sterile 0.2 μM aPES membrane and aliquoted into sterilized glass bottles. *Acetic acid was added as a concentrated 3M solution. This added a negligible amount of water to the final solution. **Molarity was calculated using manufacturer provided formula weights which includes hydration in the case of metal salt compounds.

| Environment | Compound | Molarity (mol/L)** | Stress |
| --- | --- | --- | --- |
| EnvA | NiCl<sub>2</sub> • 6H<sub>2</sub>O | 1.00 × 10<sup>-3</sup> | Metal (Nickel) |
| EnvA | Caffeine | 4.00 × 10<sup>-3</sup> | Organic |
| EnvA | Boric Acid, B(OH)<sub>3</sub> | 1.00 × 10<sup>-2</sup> | Other |
| EnvA | NaF | 1.00 × 10<sup>-2</sup> | Flouride |
| EnvB | Acetic Acid, CH<sub>3</sub>COOH* | 3.00 × 10<sup>-1</sup> | pH (acidity) |
| EnvB | LiCl | 4.00 × 10<sup>-2</sup> | Lithium |
| EnvB | Acetaminophen | 6.62 × 10<sup>-2</sup> | Organic |
| EnvB | NaCl | 1.00 × 10<sup>-1</sup> | Osmotic |
| EnvC | CrK(SO<sub>4</sub>)<sub>2</sub> • 12H<sub>2</sub>O | 2.50 × 10<sup>-3</sup> | Metal (Chromium) |
| EnvC | CuSO<sub>4</sub> • 5H<sub>2</sub>O | 2.53 × 10<sup>-2</sup> | Metal (Copper) |
| EnvC | Sodium Benzoate, C<sub>6</sub>H<sub>5</sub>COONa | 3.75 × 10<sup>-3</sup> | Benzoate |
| EnvC | ZnCl<sub>2</sub> Anhydrous | 1.25 × 10<sup>-3</sup> | Metal (Zinc) |

**Table S2.** Table of values from Figure 3. These values show the mean relative growth rates for the adapting populations in each environment; Assay 1 corresponds to Fig. 4A and Assay 2 corresponds to Fig. 4B.

| Treatment | Env A | EnvB | EnvC |
| --- | --- | --- | --- |
| A-adapted<br>(Assay 1) | 0.103 | 0.0203 | 0.0108 |
| B-adapted<br>(Assay 1) | 0.0848 | 0.135 | -0.00114 |
| C-adapted<br>(Assay 1) | 0.0280 | 0.0631 | 0.0140 |
| A-adapted<br>(Assay 2) | 0.0235 | 0.00655 | NA |
| B-adapted<br>(Assay 2) | -0.0128 | 0.0783 | NA |

**Table S3.** Table of the values from Figure 4. These values represent means and variances of relative growth rate of mutagenized genotypes, corresponding to Figure 5.

|  | A-evolved in<br>EnvA | A-evolved in<br>EnvB | B-evolved in<br>EnvA | B-evolved in<br>EnvB |
| --- | --- | --- | --- | --- |
| Mean | $-2.22 \times 10^{-3}$ | $-1.50 \times 10^{-2}$ | $-2.55 \times 10^{-2}$ | $-2.34 \times 10^{-2}$ |
| Variance | $8.89 \times 10^{-4}$ | $8.26 \times 10^{-4}$ | $6.34 \times 10^{-4}$ | $1.80 \times 10^{-3}$ |

